# All is fish that comes to the net: metabarcoding for rapid fisheries catch assessment

**DOI:** 10.1101/2020.06.18.159830

**Authors:** Tommaso Russo, Giulia Maiello, Lorenzo Talarico, Charles Baillie, Giuliano Colosimo, Lorenzo D’Andrea, Federico Di Maio, Fabio Fiorentino, Simone Franceschini, Germana Garofalo, Danilo Scannella, Stefano Cataudella, Stefano Mariani

**Affiliations:** University of Rome Tor Vergata, Department of Biology, Rome (Italy); University of Salford, School of Environment and Life Sciences, Salford (UK); San Diego Zoo, Institute for Conservation Research, San Diego (California, USA); IRBIM CNR; Liverpool John Moores University, School of Biological & Environmental Sciences, Liverpool (UK)

**Keywords:** Marine fisheries, trawling, DNA-metabarcoding, marine biodiversity, environmental impacts, eDNA

## Abstract

Monitoring marine resource exploitation is a key activity in fisheries science and biodiversity conservation. Since research surveys are time-consuming and costly, fishery-dependent data (i.e. derived directly from fishing vessels) are increasingly credited with a key role in expanding the reach of ocean monitoring. Fishing vessels may be seen as widely ranging data-collecting platforms, which could act as a fleet of sentinels for monitoring marine life, in particular exploited stocks. Here, we investigate the possibility of assessing catch composition of single hauls carried out by trawlers by applying DNA metabarcoding to the “slush” collected from fishing nets just after the end of hauling operations. We assess the performance of this approach in portraying β-diversity and examining the quantitative relationship between species abundances in the catch and DNA amount in the slush (reads counts generated by amplicon sequencing). We demonstrate that the assemblages identified using DNA in the slush mirror those returned by visual inspection of net content and detect a strong relationship between read counts and species abundances in the catch. We therefore argue that this approach could be upscaled to serve as a powerful source of information on the structure of demersal assemblages and the impact of fisheries.

## Introduction

Monitoring exploitation of marine resources and assessing the status of marine stocks and communities are key activities in fisheries science and biodiversity conservation. Achieving effective fisheries management is increasingly important as overfishing threatens fish stocks globally, reduces biodiversity, alters ecosystem functioning and jeopardizes food security and livelihoods of hundreds of millions of people worldwide (FAO 2020). Monitoring fishing activities and their impacts rely on two main sources of information: catch data (fishery-dependent) and research surveys (fishery-independent) (Dennis et al. 2015).

Since fisheries are key agents of disturbance for marine ecosystems, management requires as accurate as possible data about what, where and how much of key species is caught. The lack of such reliable catch data could lead to uncertainty about stock status, impairing our perception of resource availability, and increasing the chances of overfishing.

The collection of fishery-dependent data is historically carried-out by logbook – which are often inaccurate (Sampson 2011) – on-board observers, or self-sampling of catch done by fishers (Kraan et al. 2013). Unfortunately, several issues often prevent exhaustive data collection on catch composition. These include: discarding of non-commercial species, vessel size, distribution and operational range of the fleet, and the fact that the analysis of catch is based on time-consuming procedures such as visual sorting, taxonomic classification, counting, measuring, weighing, and/or tissue sampling. For all these reasons, the collection of fishery-dependent data is often limited to subsets of the fleet, compromising the accuracy and representativeness of the results achieved (Vilas et al. 2019). On the other hand, fishery-independent data, which are mainly collected by scientific surveys explicitly designed to capture resource distribution and status, depend on huge operational ship-time costs (Dennis et al. 2015).

Overall, collection of fishery-dependent and/or -independent data is therefore complex, time-consuming and costly. Yet, our understanding of distributions of the thousands of species caught by fisheries across the world’s oceans remains incomplete (Seebens et al. 2016), especially in developing countries (Worm and Branch 2012, Pauly and Zeller 2016), while sustainability targets, in the face of increasing climatic instability, still require spatially and temporally accurate, widespread, and affordable monitoring approaches (Bradley et al. 2019).

The rise of new technologies for the collection, management and analysis of fishery-dependent data is providing a suite of possible solutions to update and modernize fisheries data collection systems and greatly expand data collection and analysis (Bradley et al. 2019, Plet-Hansen et al. 2019). In this context, DNA metabarcoding is one major innovation that is revolutionising the way we assess biological diversity, by rapidly generating vast species inventories from trace DNA retrieved from water samples (Thomsen et al. 2012, Djurhuus et al. 2018, Stat et al. 2019), sediment cores (Fonseca et al. 2010), and bulk DNA from animal organs and tissues (McInnes et al. 2017). Different environmental media (water, sediment, gastric contents, etc.) exhibit varying levels of DNA concentration, affecting sampling, laboratory and data interpretation phases (Siegenthaler et al. 2019). However, some fishing gears, such as the nets used for bottom trawling, capture and concentrate in the net cod-end a large number of specimens in a reduced water volume. Since the amount of cells and tissues in a given volume of water should be directly related to the ratio between animal abundance and volume of water, the high biomass concentration conditions generated by trawling should lead to much greater DNA concentration of target species compared to the highly diluted conditions in seawater, commonly sampled for environmental DNA studies. This “surplus” DNA effect is expected to be even greater when animals are pressed against each other and/or wounded, losing blood or other fluids, as it typically happens during fishing operations.

In this study, we therefore asked the following questions: 1) can the concentrated water collected from fishing nets provide a readily available DNA source? 2) can DNA metabarcoding of such collections be effectively used to reconstruct catch composition? Specifically, we extracted DNA from samples of water draining from the net cod-end, just after it was hauled on board, while suspended above the deck (Fig. 1A). This water is a naturally concentrated “slush”, containing tissue and other cell material, which we hypothesized would yield DNA data reflecting species composition in the net. We used a DNA metabarcoding approach (through next-generation amplicon sequencing) to reconstruct the species composition of several hauls across different locations in the Strait of Sicily (Mediterranean Sea, Fig. 1B) and compared them directly to concurrent assessments by on-board researchers based on visual sorting and classification. Additionally, we investigated the quantitative relationship between species abundances in the catch (number and biomass of individuals) and DNA amount in the slush (read counts generated by amplicon sequencing), as, despite the enormous potential of metabarcoding, there still is little consensus on the extent to which reads correspond to the actual abundance of species (Lamb et al. 2019).

**Figure 1:**
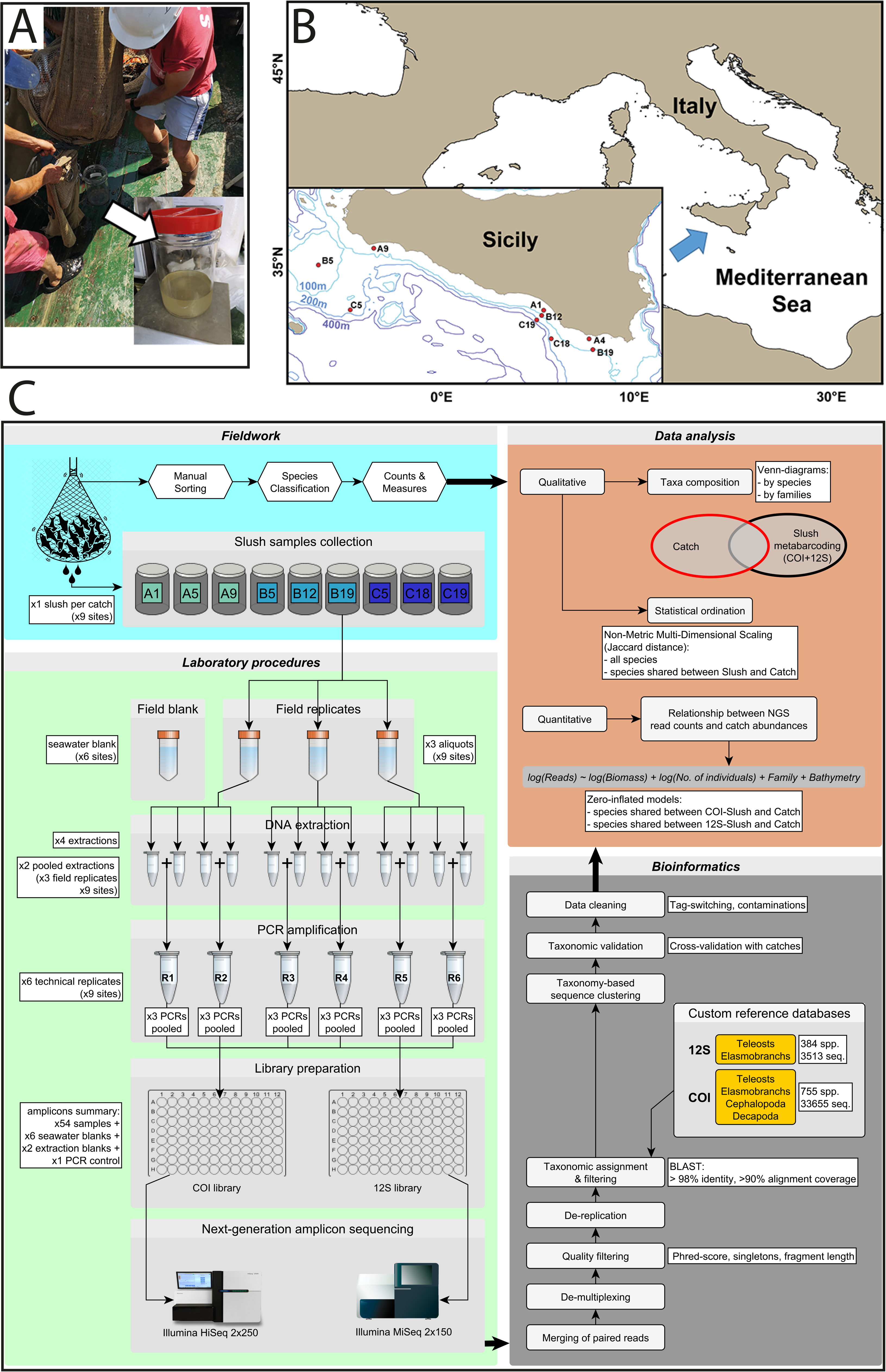
Sampling of water draining from net cod-end and study technical procedures: (A) collection of water at the end of the hauling phase; (B) Maps identifying the sampling locations along the southern coast of Sicily, Mediterranean Sea; (C) Graphical schematic illustrating the key analytical steps: fieldwork, laboratory, bioinformatics and data analyses. Colours of slush samples refer to those in Figure 3 (according to their depth stratum).

We demonstrate that the assemblages identified using DNA in the slush adequately mirror those returned by visual inspection of catch both qualitatively and quantitatively. We therefore argue that this promising approach could be upscaled to serve as a powerful source of information on the composition of demersal assemblages ad fishery impact on target stocks.

## Material and Methods

### Collection of DNA samples and catch data

Samples were obtained from nine sites (Table S1, Fig. 1B), between July and August 2018, during a bottom trawl survey within the Mediterranean International Trawl Survey (MEDITS) framework (Bertrand et al. 2002). Sampling sites covered three depth layers (10-50m, 51-100m and 101-200m). For each site, water was collected in triplicate from the dripping net cod-end just after it was hauled on board (hereafter referred to as “slush”), while suspended above the deck (Fig. 1A), then stored in three 50 ml sterile tubes (i.e. field replicates; Fig. 1C). Contextually, to account for “baseline environmental DNA contamination”, seawater was sampled nearby the vessel during hauling procedures in six out of nine sites (i.e. seawater blanks). All samples were first frozen at −40° C on board and successively transferred in the laboratory and stored at −20° C until DNA extraction.

In parallel, the species composition of each catch was determined by on-board processing of net content. All the individuals in the net were identified at the species level by visual inspection of external morphology and, if needed, analysis of meristic characters. For each species at each site we recorded the overall number of individuals and their total weight.

### DNA extraction, amplification, library preparation and Illumina sequencing

After centrifuging the slush tubes, the supernatant was removed, and 500μl of slush pellet was collected, 100μl at a time, and lysed for two hours in a ThermoMixer (Eppendorf) at 50°C and 1500 rpm, with the lysis solution from the Mu-DNA (Sellers et al. 2018) protocol for tissue. Following lysis, the rest of the protocol followed the Mu-DNA water protocol. Using a mix of these protocols allowed for efficient lysis of a more viscous sample than water (containing many cells and organismal fluids), while also removing PCR inhibitors associated with seawater. For each slush sample, four DNA extractions were performed and extracts were pooled two by two in identical volumes to maximize DNA retrieval (Fig. 1C).

We targeted a ~167bp fragment of the mitochondrial 12S gene using the Teleo02 primers (Miya et al. 2015, Taberlet et al. 2018), and a ~313bp fragment of the COI gene using the Leray-XT primers (Wangensteen et al. 2018). To uniquely distinguish each sample and aid in the detection of PCR/sequencing cross-contamination, each primer pair carried unique 8bp tags - the same tag for both forward and reverse primers. DNA extracts were PCR-amplified three times independently to minimize stochasticity. Each reaction (25μl) contained 16μl Amplitaq Gold Master Mix (Applied Biosystems), 0.16μl of Bovine Serum Albumin, 5.84μl of water, 2μl of purified DNA extract, and 1μl of each forward and reverse primer. The cycling profile for 12S primers included polymerase activation at 95°C for 10’, followed by 35 cycles of denaturation and amplification (95°C for 30”, 54°C for 45”, 72°C for 30”), and a final elongation of 72°C for 5’. The cycling profile for COI primers included polymerase activation at 95°C for 10’, followed by 35 cycles of denaturation and amplification (94°C for 1’, 45°C for 1’, 72°C for 1’), and a final elongation of 72°C for 5’. PCR replicates were then pooled prior to sequencing. Overall, we had six replicates per sites (a combination of both field and technical replicates; Fig. 1C). Note that both field and laboratory controls (i.e. six seawater blanks, two extraction blanks and a negative PCR control for each marker) were amplified to ensure quality of procedures and to assess contaminations at each step.

Amplicons were pooled and then purified with 1X paramagnetic beads (MagBio). For 12S we prepared a PCR-free, single-indexed library with the KAPA Hyper Prep Kit, following the manufacturer’s instructions. The COI library was prepared using UDI (unique dual index) adapters as it was pooled with other libraries from other projects. Libraries were cleaned of adapters and quantified using qPCR. The 12S library was loaded onto an Illumina MiSeq platform, at 8pM concentration, for 2×150 paired-end sequencing. The COI library was loaded on a larger Illumina HiSeq 2500 run, alongside samples for unrelated projects, at Macrogen Inc., for 2×250 paired-end sequencing.

### Data pre-processing

Bioinformatic procedures for quality check and data pre-processing consisted of the following steps implemented in OBITools packages (Boyer et al. 2016). First, we merged paired ends reads (minimum score = 40) and de-multiplexed samples based on their individual tag, allowing for a single base mismatch error in each tag. Second, we discarded sequences with low base-call accuracy (i.e. average quality <30 Phred score), sequences out of the expected length range (i.e. 12S = 129-209bp; COI = 303-323bp), and singletons. Third, identical sequences were collapsed (de-replication) before taxonomic assignment.

### Reference database

A reference database was created for 12S and COI, separately (Fig. 1C): the former included 12S sequences of Mediterranean fish species, while the latter contained COI sequences of Mediterranean taxa targeted by bottom trawl fishery (i.e. teleosts, elasmobranchs, cephalopods and decapods). Sequences were downloaded from the NCBI nucleotide database on 9th January 2020: 3,513 and 33,655 accessions were gathered for 12S and COI respectively. Note that the list of Mediterranean fish species (N=755) is accessible in FishBase^1^. The list of cephalopods and decapods (N=384) was obtained combining lists from SeaLifeBase^2^ and the Italian Society of Marine Biology checklists.

### Molecular taxonomic identification

We performed the taxonomic assignment of sequences that passed the previous filtering steps: unique sequences were compared to custom-made reference databases using the megaBLAST alignment algorithm implemented in the R-package rBLAST. For each query, the best match was chosen according to the maximum (bit) score. The use of such customized database i) allowed the automatic removal of non-target taxa (e.g. DNA of humans, chicken, cow) which are frequently found in such experiments due to contamination (Taberlet et al. 2018); ii) mitigated erroneous taxonomic assignments (e.g. assignment to non-Mediterranean species), which may occur when barcode sequences of a Mediterranean species are missing or shared with a non-Mediterranean species, or the barcode diversity is underrepresented compared to that of closely related exotic species. A manual inspection was performed to validate the taxonomic assignment of the more common sequences: when a sequence was shared between multiple species, we conservatively chose the one observed in catches. Only sequences showing >98% identity match combined with a >90% alignment coverage were unambiguously assigned to a species. We filtered out residual artefacts – likely originated by tag jumping (Schnell and Bohmann 2015) and/or cross-contamination – taking advantage of negative controls (i.e. six seawater blanks, two extraction blanks and a PCR negative). Specifically, the maximum frequency of reads per-species observed in negative controls (i.e. 0.11% and 0.04% for 12S and COI, respectively) was assumed as the contamination threshold. Thus, species within a sample occurring with a relative abundance below such threshold were discarded. The above-mentioned procedures are summarized in the “bioinformatics” box in Figure 1C.

### Qualitative comparison between catch and metabarcoding

To effectively compare between overall compositions of slush (data from 12S and COI were qualitatively combined together) and catch data, we drew Venn diagrams for each taxon (teleosts, elasmobranchs, cephalopods and decapods) at both species and family level. Secondly, we used non-metric multidimensional scaling (NMDS) based on Jaccard distances to assess and visualize qualitative differences in species assemblages among sampling sites and sources (i.e. visual catch examination vs. slush metabarcoding) simultaneously. NMDS was implemented in the R-package “vegan” (Oksanen et al. 2018) and performed twice including: 1) all detected species; 2) only species shared between slush and catch. In both cases, we combined COI and 12S data and replicates were treated separately.

Finally, we formally tested differences between samples, considering their source (slush or catch) as a factor of the analysis together with sampling sites, with PERMANOVA (1,000 permutations) using the vegan function “adonis”.

### Quantitative relationship between catch composition and number of DNA reads

We summed read counts from six replicates per species and site, for 12S and COI separately. Summed reads were used to investigate the quantitative relationship with catch composition, along with environmental and “taxonomic” variables, by fitting the following model:

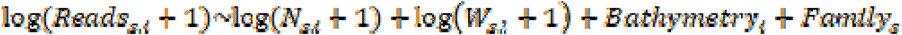

Where *Reads _s,i_* is the number of reads for the species *s* at site *i*, *N _s,i_* is the number of individuals of species *s* at site *i* (from catch data), *W _s,i_* is the weight of species *s* at site *i* (from catch data), *Bathymetry _i_* is the sea bottom depth at site *i*, and *Family _s_* is the taxonomic family of species *s*. We decided to include *Bathymetry* and *Family* variables because they both may affect the DNA amount in the slush: fishes caught in deeper sites may release more fluids than those caught in shallow water (e.g. because swim bladder often collapses during hauling procedure); while taxonomic groups may shed DNA differently, due to their physical features (e.g. crustaceans vs. molluscs). Given that the dependent variable (*Reads_s,i_*) was clearly zero-inflated (Martin et al. 2005), we fitted a zero-inflated regression model for count data via maximum likelihood, using the function “zeroinfl” of the R-package “pscl” (Jackman et al. 2020). This kind of modelling approach returns a two-component mixture model combining a binary outcome model (i.e. Bernoulli), devised to model the inflation of zero in the observed values, and a truncated count model (e.g. Poisson or negative binomial). Using this approach, it is possible to estimate the probability that a species is present and then, given it is present, estimate the relative mean number of individuals (Martin et al. 2005). Coefficients for the zero and count components, respectively, are returned.

These analyses were carried out considering only species detectable by both methods (i.e. metabarcoding and catches) to avoid that technical issues may affect the relationship (e.g. the incompleteness of reference database for taxonomic assignment). Furthermore, we removed co-absences (i.e. species missing in both slush and catches in a certain site), which would artificially improve the robustness of models. The results of this modelling exercise were represented as scatterplot overlapping a representation of the confusion matrixes generated by the comparison between expected (from catch) and observed (in the slush) number of reads per species.

## Results

After bioinformatic analysis, 12S PCR products yielded 5,433,845 reads and allowed detection of 32 species of teleosts and 8 elasmobranchs. From COI PCR products, we obtained 716,091 reads, returning 49 species of teleosts, 9 elasmobranchs, 14 cephalopods, and 5 crustaceans. Twenty-three teleosts and 4 elasmobranchs were shared between 12S and COI PCR products. Fifty-eight species were shared across slush and catch sampling methods (Fig. 2A). Over 30% of reads included the most important target species for Mediterranean demersal catches (Russo et al. 2019): the European hake (*Merluccius merluccius*), the two red mullets (*Mullus barbatus* and *Mullus surmuletus*), three species of sea bream (*Pagellus acarne*, *Pagellus bogaraveo*, and *Pagellus erythrinus*), and the deep-water rose shrimp (*Parapenaeus longirostris*). However, 30 species were detected only in the slush and 36 only in the catch. Figure 2B shows the comparison between sampling methods at the family level.

**Figure 2:**
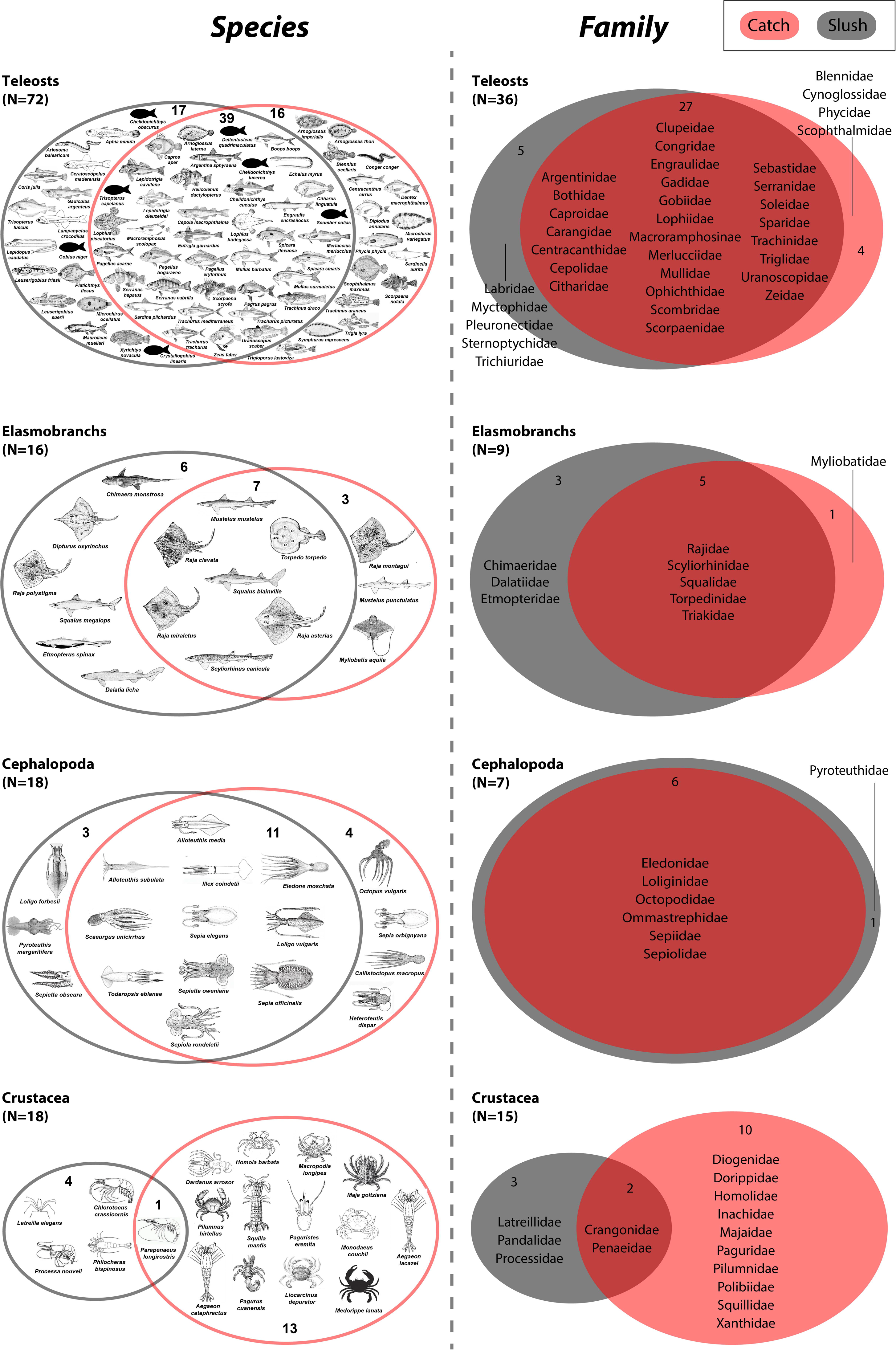
Venn diagram of species and families detected via metabarcoding of DNA in the slush and catch. Taxa identified using the 12S marker and CO1 markers are combined. Drawings were reproduced with permission of FAO Copyright Office (FAO 2017), with some exception (see Table S3).

Beta diversity reconstructed through different methods (DNA metabarcoding or visual inspection), showed a clear intra-sites affinity and a coherent distribution of samples according to the depth strata (Fig. 3). Both NMDS performed on the whole dataset (Fig. 3A) and on the subset of species shared across slush and catch (Fig. 3B) separated samples according to their spatial origin (habitat/biotope), with a clear depth gradient along the first dimension. Most notably, DNA data appear to convey very effectively the greater α- and β-diversity of the mid-depth layer (stations “B”), particularly highlighted by the divergent site B5, situated in the highly biodiverse off-shore shallow of the Adventure bank (Consoli et al. 2016).

**Figure 3:**
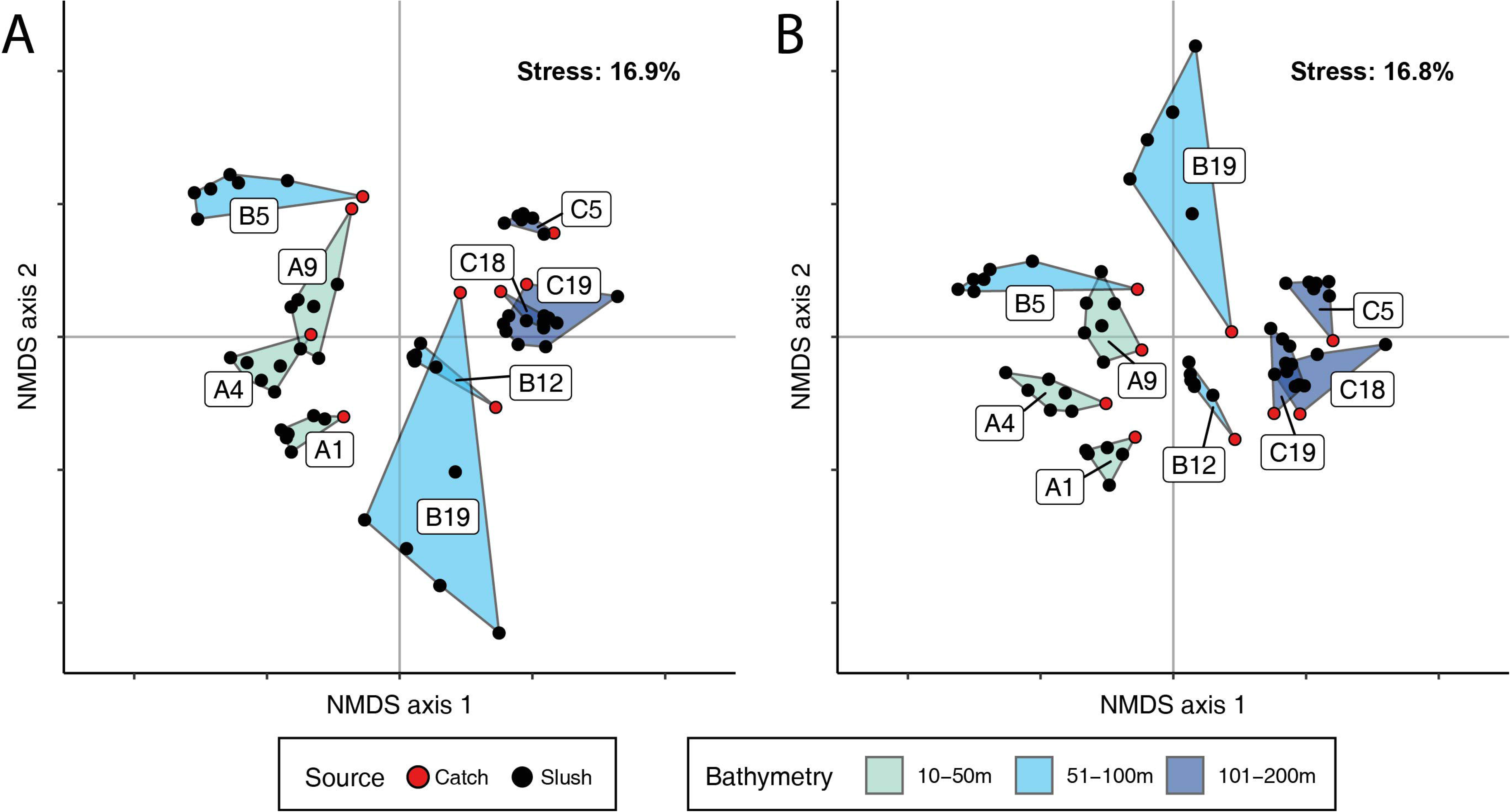
Pattern of species composition from visual sorting of survey catches and DNA sequences (12S and COI) retrieved from slush samples (six replicates per site), as returned by a Multidimensional scaling (NMDS) based on Jaccard’s distance. Samples are grouped by sampling locations into convex hulls coloured according to the corresponding depth layer and labelled according to the sampling station. A) NMDS performed on the whole dataset (125 species); B) NMDS performed on restricted dataset (58 species present in both slush and catch).

PERMANOVA on the whole dataset detected a significant difference between samples, with most of the variance explained by stations (69%), and only 3% contribution of source (i.e. slush or catch). Figure 4 shows the zero-inflated model fits (see also Table S2), in which observed values are plotted against predicted values. The zero-inflated models for 12S and COI datasets returned good fits for both zero and count components, with a clear linear relationship between predicted and observed values when both are greater than zero.

**Figure 4:**
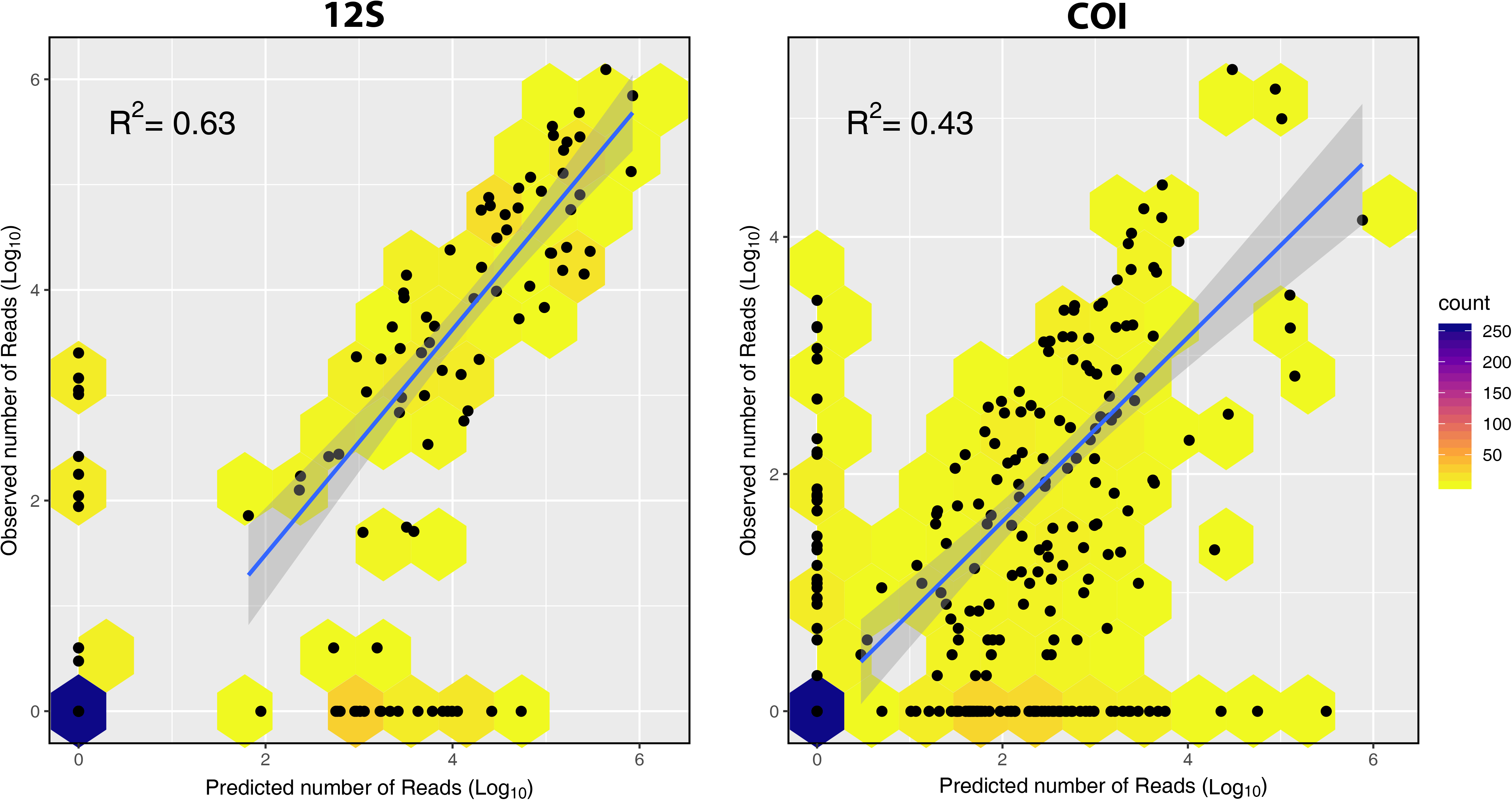
Scatterplot of observed (in the slush) and predicted (by the fitted zero-inflated models) Log_10_-transformed number of reads by species and site, for 12S and COI datasets. Points density is represented by hexagons coloured according to the scale on the right side of each plot. Linear regressions (with standard errors and corresponding adjusted R^2^ values), fitted on positive values, are also presented.

## Discussion

The idea behind this work is that fishing nets can concentrate shed material from captured species, increasing the amount of their DNA relative to the trace DNA from all the species present in the surrounding environment. Our results confirm that the water draining from trawling gear is indeed an effective source of concentrated DNA. Given that capturing sufficient quantities of DNA is one of the first critical steps in the application of eDNA-related techniques (Spens et al. 2017), and that the oceans remain largely under-surveyed, this finding opens the door to potentially important applications for fisheries and biodiversity monitoring.

Visual and DNA-based assessment methods (catch and slush) corroborate each other, indicating that water draining from net cod-end is a significantly good proxy for composition of commercial catch (Fig. 2A and Fig. 3A), and that important information on β-diversity of fishing grounds can be readily gleaned through DNA monitoring. The 30 species detected by DNA but not recovered via visual inspection reflect the power of metabarcoding approaches to identify rare and cryptic species (e.g. *R. polystigma* vs. *R. montagui*), and record taxa that are present in the environment but not catchable via trawling. Although it may appear counter-intuitive, positive detection of taxa not captured by the net can originate from a variety of processes: the physical action of the trawling gear on the substrate can suspend and retain biological material from organisms that may not be caught; this may include gametes, larvae and mucus and hence be visually undetectable in the haul; additional DNA may originate from faeces and regurgitates from the fish that are caught, and damaged, in the net. On the other hand, the 36 species that were visually identified but not detected through DNA mostly reflect the incompleteness of sequence repositories, which remains a significant challenge that must be met in the coming years in order to make DNA monitoring fully operational. This explanation is supported by the analysis at the family level, which shows a reduction of the mismatches between methods. The DNA approach appears less efficient in the case of invertebrates (crustaceans and cephalopods), and this may be partly explained by the lower sequencing depth available for COI; furthermore, as we specifically extracted DNA from centrifuged pellets (i.e. excluding the supernatant), this may have high-graded vertebrate DNA through the concentration of fish mucus.

Despite the complexities discussed above, we were able to unveil a robust correlation between number of sequence reads and species abundance in the catch. Previous studies had also suggested that metabarcoding data could be used to infer abundance, at least to some extent, and at a higher taxonomic level than the species (Thomsen et al. 2016). Here we showed that by adding few sampling-associated predictor variables to a regression model, it is possible to explain up to 63% of the variance in reads abundance across samples. Undoubtedly, the unnatural biomass concentration achieved through trawling likely underlies the more pronounced quantitative nature of metabarcoding data compared to what is normally expected in “typical” eDNA studies based on seawater samples. This represents a promising feature of the “slush” approach, for instance, in the context of catch composition reconstruction, which may often differ from recorded landings as a consequence of by-catch discard, a widespread practice that interferes with fisheries management and has substantial consequences on some stocks and assemblages.

It should be stressed that scalability of DNA-based trawl assessments would be practically achievable in a context of low-tech, rapid, non-sterile sampling operations, as fishermen and observers would not have the time to carry out sampling following strict eDNA protocols; for instance, the presence of “carry-over” DNA traces from a previous haul may still be present on the deck and the fishing gear and potentially cause noise through false positive detections in the subsequent sample site. Nevertheless, our results demonstrate that this source of bias is likely negligible: after filtering for basic contamination controls, we find that metabarcoding data mirror visual identifications, and the overall pattern of β-diversity reliably discriminates between hauls and depth strata, including sites with notable ecological features, such as the highly diverse bank at site B5.

The collection of slush samples is easy and quick (Fig. 1A), potentially providing a huge amount of data that could be collected by commercial fishing vessels operating across seas. Fishers are often the first to notice changes in marine assemblages (Bradley et al. 2019), so fishing vessels, combined with cutting-edge technology (including satellite-based tracking and electronic monitoring) are increasingly touted as potential scientific platforms not only for collecting necessary data for stock assessment but also for biodiversity data recording. The present study suggests that slush collection and storage would be a valuable low-effort task that could be carried out by most trawlers, across vast marine areas and in different seasons. This approach could be used to investigate species distribution across the oceans and, consequently, to assess species richness patterns, the spread of invasive species, and the loss of threatened and endangered species due to environmental change, ultimately providing a new tool to detect shifts in community composition (Jerde and Mahon 2015). If judiciously coordinated, fishing vessels could form an unparalleled fleet of sentinels for monitoring marine life and their changes in response to local and global perturbations, with the added bonus of potentially engendering a greater sense of marine stewardship among the fishers.

## Acknowledgements and authors contributions

TR, SM and SC conceived and designed the study; FDM, FF, GG and DS contributed to the general idea, draft the design of the sampling, and did fieldwork; FDM rinsed the samples; CB and GC performed laboratory experiments; TR, LT, GM, LDA and SF analysed data; TR, SM, LT and GM wrote the paper; all authors contributed revising the manuscript.

## Footnotes

1 https://www.fishbase.se/trophiceco/FishEcoList.php?ve_code=13

2 https://www.sealifebase.ca/speciesgroup/index.php?group=mollusks&c_code=380&action=list; https://www.sealifebase.ca/speciesgroup/index.php?group=crustaceans&c_code=380&action=list

