## Supplementary material for "All is fish that comes to the net: metabarcoding for rapid fisheries catch assessment": Table_S1

***Supplemetary Table S1***

Date, geographical coordinates (datum = WGS84), mean bathymetry, haul duration and the swept area are given for each sampling site (their codes refer to those used within the MEDITS survey framework).

| Site | Date | Lat  (°N) | Lon  (°E) | Bathymetry  (m) | Duration  (min) | Swept area  (km^2^) |
| --- | --- | --- | --- | --- | --- | --- |
| C5 | 01/08/18 | 36.951 | 12.500 | 120.5 | 30 | 0.0485 |
| B5 | 08/08/18 | 37.369 | 12.203 | 67.0 | 30 | 0.0457 |
| C19 | 10/08/18 | 36.857 | 14.237 | 169 | 30 | 0.0497 |
| B12 | 10/08/18 | 36.899 | 14.283 | 74.5 | 30 | 0.0480 |
| A1 | 10/08/18 | 36.947 | 14.302 | 21.0 | 30 | 0.0450 |
| B19 | 11/08/18 | 36.580 | 14.760 | 88.5 | 30 | 0.0457 |
| A4 | 11/08/18 | 36.680 | 14.725 | 33.5 | 30 | 0.0429 |
| C18 | 11/08/18 | 36.682 | 14.372 | 175.5 | 30 | 0.0505 |
| A9 | 16/08/18 | 37.525 | 12.719 | 42.0 | 30 | 0.0470 |
