## Supplementary material for "All is fish that comes to the net: metabarcoding for rapid fisheries catch assessment": Table_S2

***Supplementary Table S2.***

Count model coefficients and zero-inflations model coefficients for 12S and for COI. Statistically significant values (p ≤0.05) are marked in bold.

| **Model for 12S reads** | | | | |
| --- | --- | --- | --- | --- |
| **Count model coefficients (negative binomial with log link)** | | | | |
|  | **Estimate** | **Std. Error** | **z value** | **Pr(>\|z\|)** |
| *log(N + 1)* | -0.758 | 0.357 | -2.124 | **0.034** |
| *log(W + 1)* | -0.092 | 0.114 | -0.807 | 0.419 |
| *Bathymetry* | -0.005 | 0.002 | -1.887 | 0.059 |
| Caproidae | 1.442 | 0.699 | 2.064 | **0.039** |
| Carangidae | 0.797 | 0.550 | 1.450 | 0.147 |
| Clupeidae | 0.487 | 0.579 | 0.840 | 0.401 |
| Engraulidae | 0.360 | 0.512 | 0.703 | 0.482 |
| Lophiidae | 1.299 | 0.539 | 2.409 | **0.016** |
| Macroramphosinae | 3.349 | 3.004 | 1.115 | 0.265 |
| Merlucciidae | 1.030 | 0.777 | 1.325 | 0.185 |
| Ophichthidae | 1.503 | 0.591 | 2.542 | **0.011** |
| Rajidae | 1.654 | 0.462 | 3.582 | **0.000** |
| Scombridae | 0.880 | 0.526 | 1.673 | 0.094 |
| Scyliorhinidae | 1.213 | 0.614 | 1.975 | **0.048** |
| Sebastidae | 2.361 | 0.928 | 2.545 | **0.011** |
| Sparidae | 1.907 | 0.392 | 4.870 | **0.000** |
| Trachinidae | 0.711 | 0.500 | 1.423 | 0.155 |
| Triakidae | 2.108 | 0.802 | 2.629 | **0.009** |
| Zeidae | 1.744 | 0.642 | 2.716 | **0.007** |
| **Zero-inflation model coefficients (binomial with probit link)** | | | | |
|  | **Estimate** | **Std. Error** | **z value** | **Pr(>\|z\|)** |
| *log(N + 1)* | 0.505 | 0.221 | 2.284 | **0.022** |
| *log(W + 1)* | 0.264 | 0.143 | 1.843 | 0.065 |
| *Bathymetry* | 0.011 | 0.004 | 2.898 | **0.004** |
| Caproidae | 4.333 | 1.108 | 3.911 | **0.000** |
| Carangidae | 6.651 | 0.622 | 10.701 | **0.000** |
| Clupeidae | 6.087 | 0.706 | 8.628 | **0.000** |
| Engraulidae | 7.600 | 0.553 | 13.732 | **0.000** |
| Lophiidae | 4.229 | 1.126 | 3.756 | **0.000** |
| Macroramphosinae | 4.889 | 1.301 | 3.758 | **0.000** |
| Merlucciidae | 6.560 | 0.826 | 7.940 | **0.000** |
| Ophichthidae | 9.517 | 0.992 | 9.593 | **0.000** |
| Rajidae | 4.715 | 0.787 | 5.989 | **0.000** |
| Scombridae | 5.013 | 0.717 | 6.989 | **0.000** |
| Scyliorhinidae | 1.628 | 1.038 | 1.568 | 0.117 |
| Sebastidae | 3.203 | 1.546 | 2.072 | **0.038** |
| Sparidae | 6.876 | 0.658 | 10.454 | **0.000** |
| Trachinidae | 5.436 | 0.637 | 8.533 | **0.000** |
| Triakidae | 0.668 | 1.624 | 0.412 | 0.681 |
| Zeidae | 4.384 | 1.166 | 3.760 | **0.000** |
| Log(theta) | -0.466 | 0.148 | -3.156 | **0.002** |

| **Model for COI reads** | | | | |
| --- | --- | --- | --- | --- |
| **Count model coefficients (negative binomial with log link)** | | | | |
|  | **Estimate** | **Std. Error** | **z value** | **Pr(>\|z\|)** |
| log(N + 1) | 1.267 | 0.285 | 4.441 | **0.000** |
| log(W + 1) | -0.389 | 0.177 | -2.203 | **0.028** |
| Bathymetry | 0.000 | 0.004 | -0.025 | 0.980 |
| Argentinidae | 4.380 | 1.282 | 3.418 | **0.001** |
| Bothidae | 4.232 | 0.895 | 4.726 | **0.000** |
| Caproidae | 2.615 | 1.157 | 2.259 | **0.024** |
| Carangidae | 5.015 | 0.734 | 6.831 | **0.000** |
| Centracanthidae | 5.972 | 0.853 | 6.999 | **0.000** |
| Cepolidae | 0.812 | 1.559 | 0.521 | 0.602 |
| Citharidae | 5.470 | 1.292 | 4.235 | **0.000** |
| Clupeidae | 2.933 | 1.223 | 2.398 | **0.016** |
| Eledonidae | 4.003 | 1.194 | 3.352 | **0.001** |
| Engraulidae | 5.372 | 0.958 | 5.607 | **0.000** |
| Gadidae | 5.301 | 2.360 | 2.246 | **0.025** |
| Gobiidae | 6.435 | 1.502 | 4.285 | **0.000** |
| Loliginidae | 4.966 | 0.576 | 8.623 | **0.000** |
| Lophiidae | 3.103 | 2.823 | 1.099 | 0.272 |
| Macroramphosinae | 2.685 | 1.153 | 2.328 | **0.020** |
| Merlucciida | 4.704 | 1.001 | 4.699 | **0.000** |
| Mullidae | 6.960 | 0.888 | 7.841 | **0.000** |
| Octopodidae | 3.009 | 1.613 | 1.865 | 0.062 |
| Ommastrephidae | 6.489 | 0.978 | 6.638 | **0.000** |
| Ophichthidae | 11.661 | 1.791 | 6.509 | **0.000** |
| Penaeidae | 2.961 | 1.552 | 1.908 | 0.056 |
| Rajidae | 5.599 | 1.381 | 4.054 | **0.000** |
| Scombridae | 4.703 | 1.673 | 2.810 | **0.005** |
| Scorpaenidae | 8.533 | 2.621 | 3.255 | **0.001** |
| Scyliorhinidae | 4.473 | 1.541 | 2.903 | **0.004** |
| Sebastidae | 3.075 | 2.501 | 1.230 | 0.219 |
| Sepiidae | 4.980 | 0.827 | 6.019 | **0.000** |
| Sepiolidae | 5.459 | 1.105 | 4.941 | **0.000** |
| Serranidae | 2.442 | 0.758 | 3.222 | **0.001** |
| Sparidae | 4.550 | 0.802 | 5.674 | **0.000** |
| Squalidae | 4.275 | 2.529 | 1.690 | 0.091 |
| Torpedinidae | 5.605 | 2.456 | 2.282 | **0.022** |
| Trachinidae | 4.384 | 1.510 | 2.903 | **0.004** |
| Triglidae | 3.090 | 0.920 | 3.359 | **0.001** |
| Uranoscopidae | 2.161 | 2.472 | 0.874 | 0.382 |
| Zeidae | 3.295 | 1.217 | 2.708 | **0.007** |
| Log(theta) | -1.664 | 0.104 | -16.062 | **0.000** |
| **Zero-inflation model coefficients (binomial with probit link)** | | | | |
|  | **Estimate** | **Std. Error** | **z value** | **Pr(>\|z\|)** |
| *log(N + 1)* | 1.952 | 1.532 | 1.275 | 0.202 |
| *log(W + 1)* | -2.055 | 1.397 | -1.471 | 0.141 |
| *Bathymetry* | -0.001 | 0.003 | -0.544 | 0.587 |
| Argentinidae | 0.624 | 0.788 | 0.792 | 0.428 |
| Bothidae | -0.614 | 1.059 | -0.580 | 0.562 |
| Caproidae | 0.496 | 0.985 | 0.504 | 0.614 |
| Carangidae | 0.519 | 0.536 | 0.969 | 0.332 |
| Centracanthidae | 0.852 | 0.557 | 1.529 | 0.126 |
| Cepolidae | 1.899 | 1.453 | 1.307 | 0.191 |
| Citharidae | 3.662 | 7.383 | 0.496 | 0.620 |
| Clupeidae | 0.753 | 0.854 | 0.881 | 0.378 |
| Eledonidae | 0.310 | 0.725 | 0.428 | 0.669 |
| Engraulidae | -0.328 | 0.829 | -0.396 | 0.692 |
| Gadidae | 3.343 | 3.911 | 0.855 | 0.393 |
| Gobiidae | 0.639 | 0.600 | 1.064 | 0.287 |
| Loliginidae | -0.501 | 0.851 | -0.588 | 0.556 |
| Lophiidae | 5.220 | 213.340 | 0.024 | 0.980 |
| Macroramphosinae | 0.540 | 0.785 | 0.689 | 0.491 |
| Merlucciidae | -0.544 | 1.569 | -0.347 | 0.729 |
| Mullidae | 1.113 | 0.777 | 1.442 | 0.149 |
| Octopodidae | 3.962 | 13.466 | 0.294 | 0.769 |
| Ommastrephidae | 0.499 | 0.507 | 0.986 | 0.324 |
| Ophichthidae | 8.106 | 5.554 | 1.459 | 0.144 |
| Penaeidae | 2.796 | 1.491 | 1.876 | 0.061 |
| Rajidae | 1.649 | 0.553 | 2.982 | **0.003** |
| Scombridae | 2.603 | 1.889 | 1.377 | 0.168 |
| Scorpaenidae | 10.236 | 8.320 | 1.230 | 0.219 |
| Scyliorhinidae | 4.829 | 102.284 | 0.047 | 0.962 |
| Sebastidae | 4.847 | 65.165 | 0.074 | 0.941 |
| Sepiidae | 0.651 | 0.501 | 1.299 | 0.194 |
| Sepiolidae | 1.123 | 0.498 | 2.256 | **0.024** |
| Serranidae | 0.427 | 0.693 | 0.616 | 0.538 |
| Sparidae | 1.806 | 0.649 | 2.782 | **0.005** |
| Squalidae | 5.185 | 177.417 | 0.029 | 0.977 |
| Torpedinidae | 5.174 | 158.587 | 0.033 | 0.974 |
| Trachinidae | 1.013 | 0.681 | 1.488 | 0.137 |
| Triglidae | 0.834 | 0.446 | 1.871 | 0.061 |
| Uranoscopidae | 4.967 | 37.890 | 0.131 | 0.896 |
| Zeidae | -5.846 | NA | NA | NA |
