## Supplementary material for "All is fish that comes to the net: metabarcoding for rapid fisheries catch assessment": Table_S3

**Supplementary table S3.** Source of some drawing for Figure 2

| *Loligo subulata* | https://wikivisually.com/wiki/Alloteuthis_subulata |
| --- | --- |
| *Medorippe lanata* | https://it.freepik.com/vettori-premium/granchi-bairdi_5969456.htm#page=1&position=15 |
| *Pagurus cuanensis* | http://megabenthos.info/catalog/arthropoda/malacostraca/decapoda/paguridae/pagurus/pagurus-cuanensis/ |
| *Pagurus cuanensis* | <https://it.wikipedia.org/wiki/File:Pyroteuthismargaritifera.jpg#file> |
| *Sepietta oweniana* | https://en.wikipedia.org/wiki/File:Sepietta_oweniana_-_from_Commons.jpg |
| *Paguristes eremita* | https://decapoda.nhm.org/pdfs/21446/21446.pdf |
